# Population genomics of parallel hybrid zones in the mimetic butterflies, *H. melpomene* and *H. erato*

**DOI:** 10.1101/000208

**Authors:** Nicola J. Nadeau, Mayté Ruiz, Patricio Salazar, Brian Counterman, Jose Alejandro Medina, Humberto Ortiz-Zuazaga, Anna Morrison, W. Owen McMillan, Chris D. Jiggins, Riccardo Papa

**Affiliations:** Department of Zoology, University of Cambridge, UK; Department of Animal and Plant Sciences, University of Sheffield, UK; Department of Biology and Center for Applied Tropical Ecology and Conservation, University of Puerto Rico, Rio Piedras, San Juan, Puerto Rico 00921; Centro de Investigación en Biodiversidad y Cambio Climático (BioCamb), Universidad Tecnológica Indoamérica, Quito, Ecuador; Department of Biology, Mississippi State University, USA; High Performance Computing Facility, University of Puerto Rico, Puerto Rico; Department of Computer Science, University of Puerto Rico Rio Piedras, Puerto Rico; Smithsonian Tropical Research Institute, Panama

**Keywords:** Hybrid zones, convergent evolution, adaptive divergence, genome-wide association mapping, RAD sequencing

## Abstract

Hybrid zones can be valuable tools for studying evolution and identifying genomic regions responsible for adaptive divergence and underlying phenotypic variation. Hybrid zones between subspecies of *Heliconius* butterflies can be very narrow and are maintained by strong selection acting on colour pattern. The co-mimetic species *H. erato* and *H. melpomene* have parallel hybrid zones where both species undergo a change from one colour pattern form to another. We use restriction associated DNA sequencing to obtain several thousand genome wide sequence markers and use these to analyse patterns of population divergence across two pairs of parallel hybrid zones in Peru and Ecuador. We compare two approaches for analysis of this type of data; alignment to a reference genome and *de novo* assembly, and find that alignment gives the best results for species both closely (*H. melpomene*) and distantly (*H. erato*, ^∼^15% divergent) related to the reference sequence. Our results confirm that the colour pattern controlling loci account for the majority of divergent regions across the genome, but we also detect other divergent regions apparently unlinked to colour pattern differences. We also use association mapping to identify previously unmapped colour pattern loci, in particular the *Ro* locus. Finally, we identify within our sample a new cryptic population of *H. timareta* in Ecuador, which occurs at relatively low altitude and is mimetic with *H. melpomene malleti*.

## INTRODUCTION

Natural hybrid zones occur where divergent forms meet, mate and hybridise. Narrow hybrid zones can be maintained by strong selection that prevents mixing or favours particular forms in particular areas (Barton and Hewitt 1985). Studies of hybrid zones have provided many insights into the origins of diversity and the process of speciation (Mallet et al. 1990; Kawakami and Butlin 2001; Harrison 1993). High-throughput sequencing technologies now provide the opportunity for hybrid zones to fully meet their potential as windows into the evolutionary process, by allowing us to move beyond studies of neutral variation at a handful of loci, to describing the genetic basis of phenotypic differences and identifying loci under selection (Crawford and Nielsen 2013; Rieseberg and Buerkle 2002; Gompert et al. 2012).

Butterflies of the genus *Heliconius* are extremely diverse in their wing colour patterns, and combine extensive intra-specific diversity with perfect inter-specific similarity in wing phenotypes. The bright wing colourations that identify this group of Neotropical butterflies are used as aposematic warnings to predators, and are under positive frequency dependent selection, which favours common colour patterns that predators learn to avoid. This strong selection also maintains narrow hybrid zones between subspecies with different patterns (Benson 1972; Mallet and Barton 1989a; Kapan 2001; Langham 2004). In addition, frequency dependent selection has led to Müllerian mimicry between many distinct species (Müller 1879). For instance, *H. erato* and *H. melpomene* are two distantly related species that diverged around 15-20 million years ago, but have converged on common colour patterns across most of the Neotropics, with parallel hybrid zones between subspecies of both species (Figure 1). Current evidence suggests that the convergent colour patterns in these species have most likely evolved independently (Supple et al 2013, Hines et al. 2011). It has been suggested that *H. erato* is more ancient and *H. melpomene* diversified more recently to mimic the *H. erato* forms (Brower 1996; Flanagan et al. 2004; Quek et al. 2010). Nevertheless, it appears that the same handful of genetic loci are responsible for producing most of the colour pattern variation observed in both species (Joron et al. 2006; Baxter et al. 2008; Martin et al. 2012; Reed et al. 2011). This pattern of parallel adaptive evolution, with multiple instances of both species convergently evolving highly similar phenotypes, makes this an excellent system in which to address questions about the predictability of the evolutionary process and the extent to which particular genes are re-used when evolving the same phenotypes (Nadeau and Jiggins 2010; Papa et al. 2008).

**Figure 1.**
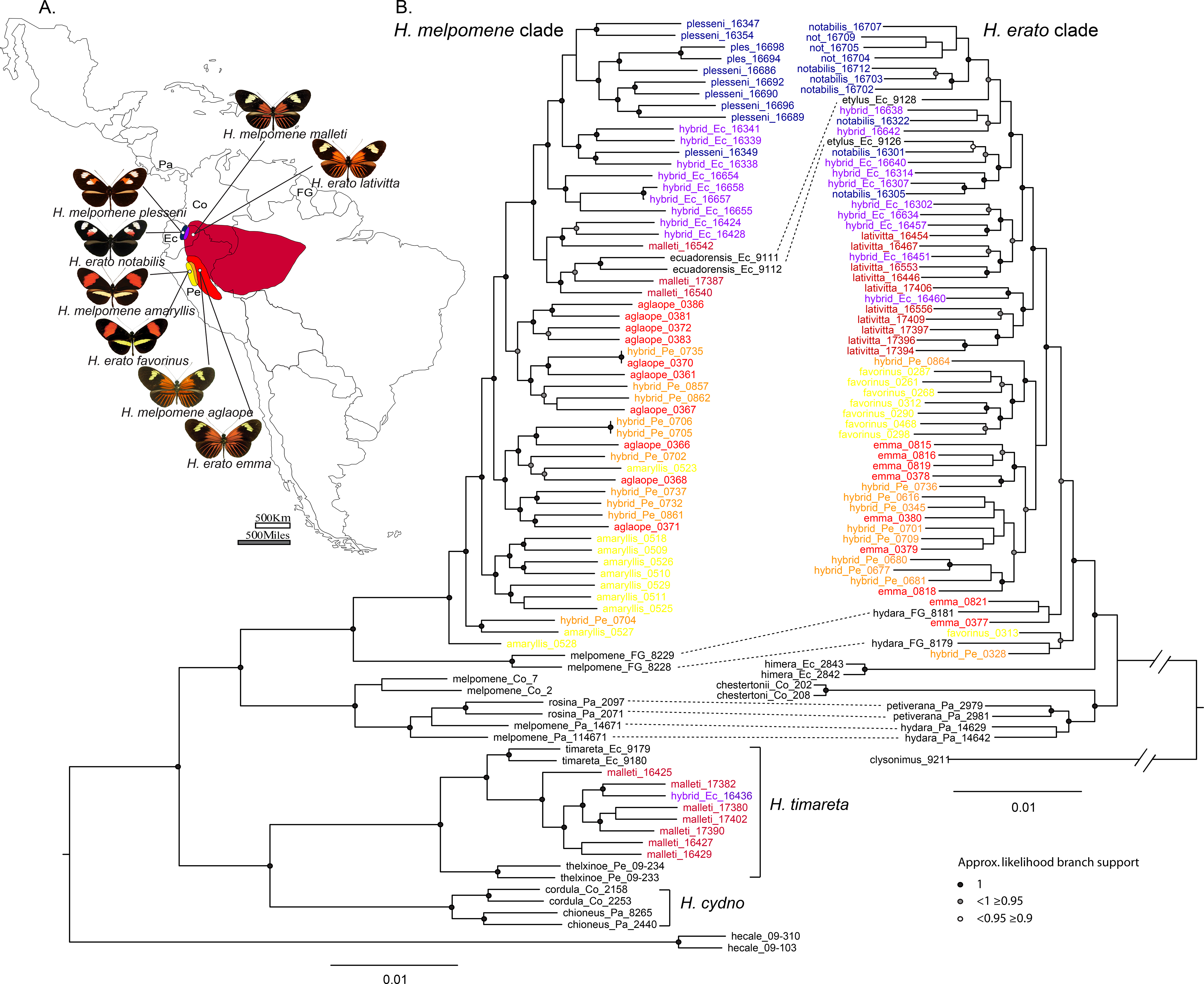
A) Distribution in South America of the subspecies included in this study. B) Maximum likelihood phylogenies with approximate likelihood branch supports. Co-mimics from outside of the focal hybrid zones are connected with dotted lines. Focal hybrid zone individuals are shown in colour: blue, *H. m. plesseni* and *H. e. notabilis*; purple, Ecuador hybrids; dark red, *H. m. malleti* and *H. e. lativitta*; red, *H. m. aglaope* and *H. e. favorinus*; orange, Peru hybrids; yellow, *H. m. amaryllis* and *H. e. emma*. Additional populations are in black. Country abbreviations: Ec, Ecuador; FG, French Guiana; Co, Colombia; Pa, Panama.

In this study, we use high resolution genome scans to investigate patterns of divergence across two pairs of parallel hybrid zones, in Peru and Ecuador. These occur between subspecies with different wing colour patterns in both *H. erato* and *H. melpomene* (Figure 1). In both regions the clines in colour pattern alleles between species are highly coincident (Mallet et al. 1990; Salazar 2012). The two hybrid zones in Peru have been the focus of several previous studies, while those in Ecuador have been less well studied. In Peru, strong natural selection has been shown to maintain colour pattern differences (Mallet and Barton 1989a) and the colour pattern controlling loci show increased divergence compared to other loci in the genome (Baxter et al. 2010; Counterman et al. 2010; Nadeau et al. 2012; Supple et al. 2013; S. H. Martin et al. 2013). However, we are lacking a clear picture of exactly how many loci are divergent between subspecies, and the extent to which the genomic architecture of divergence is the same between mimetic species.

Extensive genetic mapping using experimental crosses between different colour pattern forms has identified the chromosomal regions responsible for colour pattern variation in these species (Sheppard et al. 1985; Baxter et al. 2008; Joron et al. 2006; Papa et al. 2013). Three major clusters of loci control most of the colour pattern variation observed in both species. The tightly linked *B* and *D* loci in *H. melpomene* control the red forewing band and the red/orange hindwing rays and proximal “dennis” patches on both wings respectively. These loci are homologous to the *D* locus in *H. erato* (Baxter et al. 2008) and appear to be cis regulatory elements of the *optix* gene on chromosome 18 (Reed et al. 2011; Supple et al. 2013). The *Ac* and *Sd* loci, in *H. melpomene* and *H. erato* respectively, control the shape of the forewing band via regulation of the *WntA* gene on chromosome 10 (Martin et al. 2012). The presence of most yellow and white elements on the wing is largely controlled by three tightly linked loci, *Yb, Sb* and *N*, on chromosome 15 in *H. melpomene* (Ferguson et al. 2010), which are homologous to the *Cr* locus in *H. erato* (Joron et al. 2006). Quantitative trait locus (QTL) mapping has identified other loci of minor effect, including at least 7 additional QTL in *H. erato* (Papa et al. 2013), and QTL in *H. melpomene* on chromosomes 2, 7 and 13 that affect forewing band shape (Baxter et al. 2008). In some cases mapping studies have been followed up by population genetic studies of the mapped intervals across natural hybrid zones where many generations of crossing and backcrossing have led to narrow regions of association, permitting fine scale mapping (Baxter et al. 2010; Counterman et al. 2010; Nadeau et al. 2012; Supple et al. 2013). High-throughput sequencing technologies now provide the feasibility to generate a suitably high density of genomic markers to identify the narrow QTL present in these hybrid zones without the necessity of performing controlled laboratory crosses (Crawford and Nielsen 2013). In this study we aim to test this approach, using this system where we already know some of the loci responsible for phenotypic differences.

The hybrid zones that we focus on all occur across altitudinal gradients (Figure 2A). Therefore, it is possible that traits other than colour pattern may be differentiated across the hybrid zones driven by altitudinal selection, for example related to temperature or changes in larval host plants. Such selection on additional regions of the genome could help to stabilise the hybrid zone, which, if due to frequency dependent selection acting on colour pattern alone, could be unstable (Bierne et al. 2011; Barton and Hewitt 1985; Mallet and Barton 1989b; Mallet 2010). Therefore an important question that we will address is whether there are divergent regions of the genome that are not controlling colour pattern, as these could be candidates for loci controlling other aspects of ecological adaptation.

**Figure 2.**
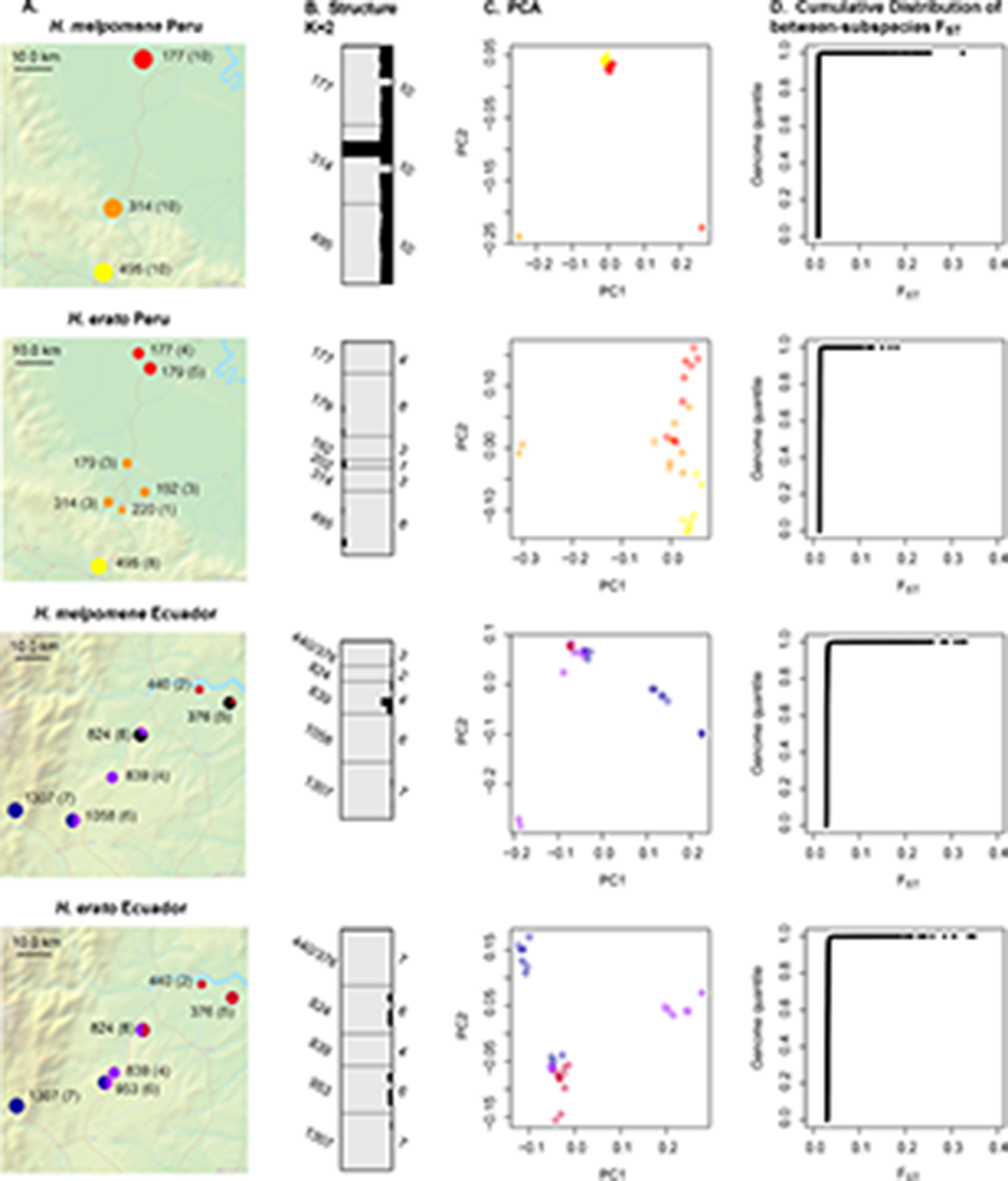
Population structure at each of the hybrid zones using the reference aligned data. A) Sampling locations with altitude (sample number) and pie charts of the proportion of individuals of each type sampled from each site. Colours are the same as in Figure 1 except black indicates *H. timareta* in Ecuador. B) Structure analysis with k=2 (*H. timareta* individuals excluded). Each individual is shown as a horizontal bar with the allelic contribution from population 1 (grey) and population 2 (black) C) Principle components analysis. D) Distribution of *F_ST_* values from BayeScan

In this study, we use restriction associated DNA (RAD) sequencing (Baird et al. 2008) to determine, for the first time:

1. if association mapping in these hybrid zones can identify known and novel loci underlying phenotypic variation
2. how much of the genome is differentiated and under divergent selection between subspecies
3. how much of this differentiation is due to loci controlling colour pattern variation
4. if the same regions are divergent between co-mimetic species

We also investigate the advantages and limitations of alignment and assembly methods when only a single reference genome is available. We compare two widely used approaches: *de novo* assembly of restriction associated reads using the program Stacks (Catchen et al. 2011) versus alignment of paired end reads to the reference *H. melpomene* genome.

## RESULTS

### Summary of the data and comparison of alignment and assembly techniques

We sequenced a total of 129 individuals of *H. erato* and *H. melpomene* from the four hybrid zones in Peru and Ecuador, and including a small number of additional individuals from across the range of *H. erato*. Using restriction associated DNA sequencing (RAD-seq), we obtained a total of 1,496M 150 base paired-end reads from the hybrid zone individuals, and an additional 115M 100 base paired-end reads from the other *H. erato* populations and outgroups. We also include in our analyses data from additional *H. melpomene* populations and outgroups already published in a previous study (Nadeau et al. 2013).

Our reference genome for *H. melpomene* is highly divergent from *H. erato*. Nonetheless, for both species, alignment of reads to the reference yielded more usable data when compared to *de novo* assembly. The latter produced more bases in assembled contigs (Table 1), but only ^∼^2% of contigs assembled in the *H. erato* populations were present in more than 10 individuals, with the figure being approximately 7% in *H. melpomene*. By comparison when the same data (plus the paired-end reads) were aligned to our reference sequence, approximately 38% of aligned bases were found in more than 10 individuals in *H. erato* and >50% in *H. melpomene*. We hypothesised that high levels of within population variation led to homologous reads being separated into distinct contigs in the *de novo* assembly. We could confirm that this was the case for one region of the *H. erato* genome for which a high quality reference sequence is available (Supple 2013). Across 960 kb at the *D* colour pattern locus, we observed that in regions with high sequence divergence between subspecies, our RAD-seq reads were assembled into separate contigs. Overall, we also found a higher frequency of single nucleotide polymorphisms (SNPs) in the reference alignments than the *de novo* assemblies (Table 1). These SNPs were defined as sites that were polymorphic within the sampled populations and so are not inflated by fixed differences from the reference genome.

**Table 1.**
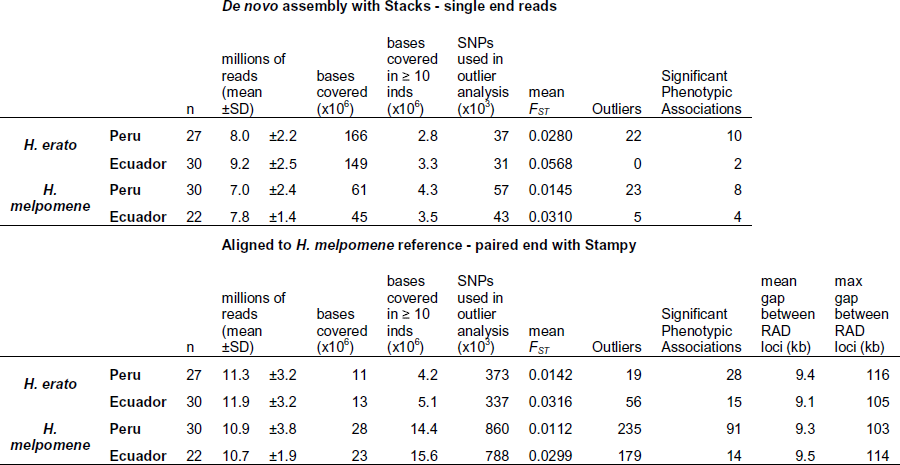
Summary statistics from alignment and assembly approaches.

| De novo assembly with Stacks - single end reads |  |  |  |  |  |  |  |  |  |  |  |  |
| --- | --- | --- | --- | --- | --- | --- | --- | --- | --- | --- | --- | --- |
|  |  | n | millions of reads (mean ±SD) |  | bases covered (x10<sup>6</sup>) | bases covered in ≥ 10 inds (x10<sup>6</sup>) | SNPs used in outlier analysis (x10<sup>3</sup>) | mean *F*<sub>ST</sub> | Outliers | Significant Phenotypic Associations |  |  |
| *H. erato* | Peru | 27 | 8.0 | ±2.2 | 166 | 2.8 | 37 | 0.0280 | 22 | 10 |  |  |
|  | Ecuador | 30 | 9.2 | ±2.5 | 149 | 3.3 | 31 | 0.0568 | 0 | 2 |  |  |
| *H. melpomene* | Peru | 30 | 7.0 | ±2.4 | 61 | 4.3 | 57 | 0.0145 | 23 | 8 |  |  |
|  | Ecuador | 22 | 7.8 | ±1.4 | 45 | 3.5 | 43 | 0.0310 | 5 | 4 |  |  |
| Aligned to *H. melpomene* reference - paired end with Stampy |  |  |  |  |  |  |  |  |  |  |  |  |
|  |  | n | millions of reads (mean ±SD) |  | bases covered (x10<sup>6</sup>) | bases covered in ≥ 10 inds (x10<sup>6</sup>) | SNPs used in outlier analysis (x10<sup>3</sup>) | mean *F*<sub>ST</sub> | Outliers | Significant Phenotypic Associations | mean gap between RAD loci (kb) | max gap between RAD loci (kb) |
| *H. erato* | Peru | 27 | 11.3 | ±3.2 | 11 | 4.2 | 373 | 0.0142 | 19 | 28 | 9.4 | 116 |
|  | Ecuador | 30 | 11.9 | ±3.2 | 13 | 5.1 | 337 | 0.0316 | 56 | 15 | 9.1 | 105 |
| *H. melpomene* | Peru | 30 | 10.9 | ±3.8 | 28 | 14.4 | 860 | 0.0112 | 235 | 91 | 9.3 | 103 |
|  | Ecuador | 22 | 10.7 | ±1.9 | 23 | 15.6 | 788 | 0.0299 | 179 | 14 | 9.5 | 114 |

As expected, given that *H. erato* is ^∼^15% divergent from *H. melpomene* in the aligned data, fewer *H. erato* reads aligned to the *H. melpomene* genome as compared to those from *H. melpomene*, leading to fewer confidently called bases. Nevertheless, the use of the reference *H. melpomene* genome for aligning the *H. erato* reads resulted in more bases being called across multiple individuals and around 10x more SNPs identified when compared to the *de novo* assembly approach. In addition, the gaps between aligned RAD loci were similar across both species (Table 1), indicating that the reduced number of bases is not due to fewer RAD loci aligning but to fewer confidently called bases at each RAD locus. The power to detect loci under selection or responsible for phenotypic variation should therefore be similar in both species. In summary, it seems that the aligned data should give the most power to detect divergent regions and phenotypic associations for both species. However, we performed outlier and association analyses using the output of both approaches for comparison. It is also possible that the *de novo* assembly might detect divergent regions important in adaptation that could not be aligned to the *H. melpomene* reference.

### Phylogenetics and population structure

Using the reference aligned sequence data, we constructed maximum likelihood phylogenies for the *H. melpomene* and *H. erato* clades, including individuals from additional populations and outgroup taxa (Figure 1). This revealed remarkably similar patterns of divergence between co-occurring, co-mimetic subspecies in both groups. Population divergence in *H. erato* is thought to be deeper than that in *H. melpomene* (Flanagan et al. 2004 but see Cuthill and Charleston 2012), but this was not evident in our tree as branch lengths were very similar between the two species. However this may be due to the poorer alignments for *H. erato*, with the *H. erato* tree based only on about a third as many sites as that for *H. melpomene*. These sites in *H. erato* are likely to be more conserved, resulting in some compression of the tree topology.

The most striking finding from the phylogenetic reconstruction was that eight of the presumed *H. melpomene* individuals from Ecuador were strongly supported as clustering within the *H. timareta* clade (Figure 1). All of these individuals had a *H. melpomene malleti-*like phenotype with the exception of one individual which had been characterised as a possible hybrid due to a large and rounded yellow forewing band, but was otherwise *H. m. malleti-*like. This finding was surprising because while populations of *H. timareta* mimetic with *H. m. malleti* have previously been described in Colombia (Giraldo et al. 2008) and Northern Peru (Lamas 1997), they are all found in highland areas above ^∼^1000 m. Similar populations are not known from lowland sites anywhere in the range. To compare our individuals to these and other populations, we also directly sequenced part of the mitochondrial COI gene that overlaps with the region sequenced in previous studies (Giraldo et al. 2008; Mérot et al. 2013). Our phylogeny based on these sequences also robustly supported these eight individuals as being *H. timareta* and placed them closer to the highland *H. timareta timareta* in Ecuador than to *H. timareta florencia* in Colombia that they resemble phenotypically (SI figure 1).

The newly identified *H. timareta* subspecies was also clearly evident in a principle components analysis (PCA) of the combined *H. melpomene, H. timareta* and *H. cydno* data. The first principle component separated the Peruvian *H. melpomene* from *H. timareta* and *H. cydno* (which were very similar on this axis, SI figure 2). The grouping of the Ecuadorian samples was consistent with the phylogeny, with the same eight individuals clustering with *H. timareta*. No individuals were intermediate between *H. melpomene* and *H. timareta*, indicating that the level of genetic isolation between the two species is similar to elsewhere in their range. This was also confirmed by a Structure analysis of the Ecuador *“H. melpomene”* population, where a model with two populations had the best fit to the data (posterior probability = 1). Under this model, which allowed for admixture between populations, the *H. timareta* individuals all had 100% of their allelic contribution from population 1, while for *H. melpomene* the maximum contribution of population 1 to any individual’s genotype was 1.8% (SI Table 1). In summary, we can conclude that these are distinct species with little gene flow between them.

We conducted further analyses of the genetic structure of each of the hybrid zone populations, excluding the *H. timareta* individuals, again using the reference aligned data. Overall, these results suggest only very low genetic differentiation between any of the parapatric subspecies. Structure analyses of each population generally showed very little structure and strongest support for only a single population cluster being present. The only exception was the Peruvian *H. melpomene*, where two population clusters gave the highest posterior probability (p = 1). However, these clusters did not correspond to the two subspecies. The genetic diversity was partitioned such that most individuals were admixed with about a quarter of their allelic variation from population 2, except for two “hybrid” individuals that had pure population 2 genotypes and two other individuals (one “hybrid” and one *aglaope*) that had almost pure population 1 genotypes (Figure 2B). PCA revealed very similar patterns, with small groups of hybrid phenotype individuals giving the clearest clusters, which in most cases were also identified by Structure (with K = 2, Figure 2C). Three of the populations did reveal some separation of the subspecies at one of the first two principle components, but with a gradual change from one genomic “type” to another. The *H. melpomene* subspecies in Ecuador were separated by PC1, which explained 10% of the variation in this population. The two *H. erato* populations both showed some separation by subspecies at PC2, which explained 5.7% and 6.7% of the variation in Peru and Ecuador respectively. We found very similar results with PCA on the *de novo* assembled data (SI figure 3), suggesting that the underlying genetic signal in both data sets is very similar. The lack of strong differentiation between subspecies was also supported by the *F_ST_* distributions (calculated by BayeScan), which gave very low *F_ST_* values between subspecies at over 99% of the genome, with only a small percentage of SNPs showing high levels of differentiation (Figure 2D, SI figure 3).

### Association mapping of loci responsible for phenotypic variation

We performed association mapping to identify genetic regions responsible for the phenotypic variation that segregates across each of the hybrid zones. In general, the expected associations were found at the three major loci known to control colour pattern variation on chromosomes 10,15 and 18 (Figures 3A/D, 4A/D and Table 2). The majority of SNPs showing significant phenotypic associations fell within or tightly linked to these loci in all populations except in Peruvian *H. erato*, where only 26% were tightly linked to the known loci.

**Figure 3.**
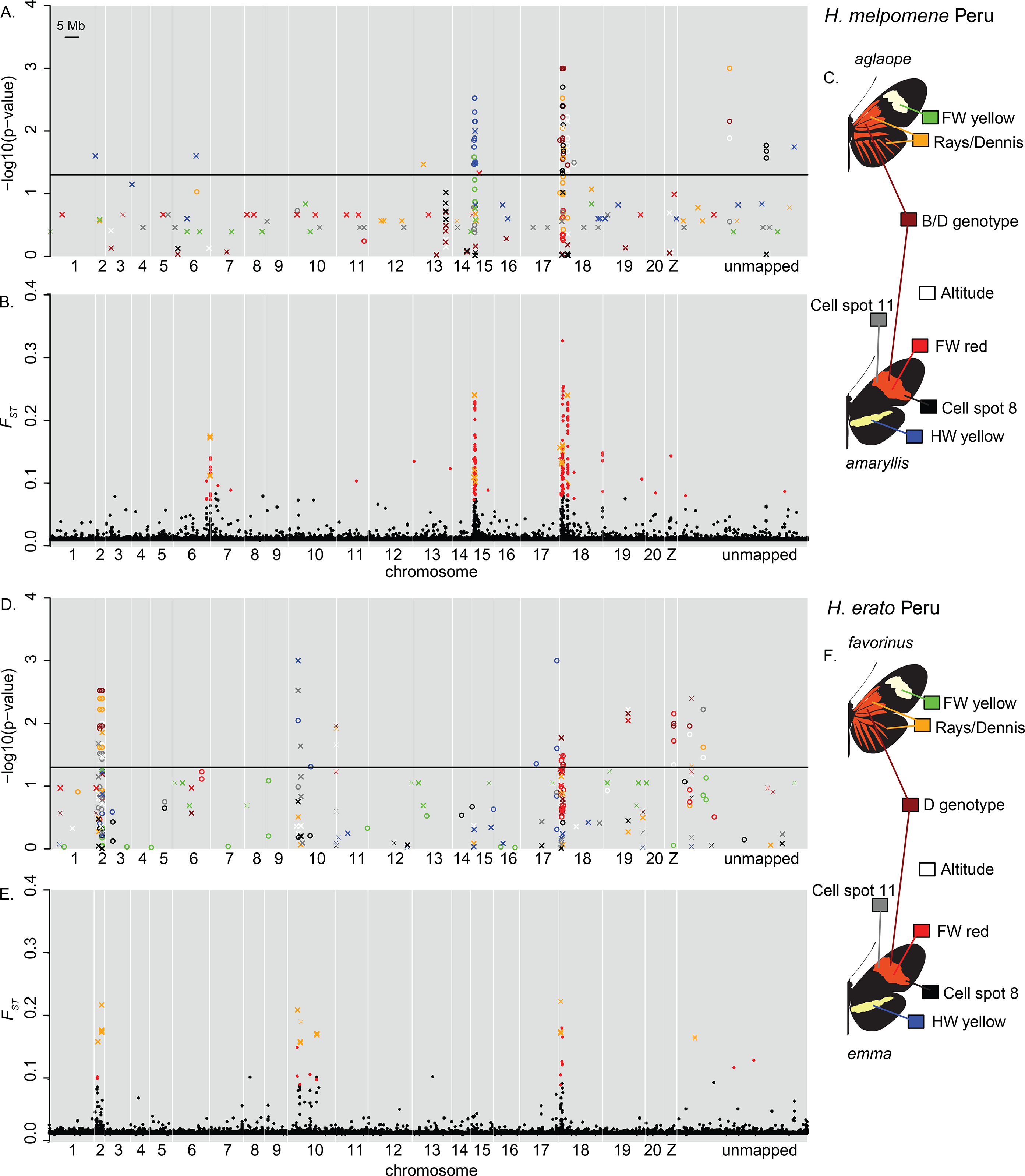
Association mapping (A and D) and outlier analysis (B and E) for *H. melpomene* (A, B, C) and *H. erato* (D, E, F) in Peru. Each phenotype used for the association mapping is shown in a different colour as illustrated in panels C and F. For clarity, only the top 20 associated SNPs are shown for each phenotype. Results from the *de novo* assembled data are shown as crosses (and in orange for the outlier analysis) and positioned based on the top BLAST hit to the *H. melpomene* genome, those with thinner lines were not confidently or uniquely assigned to these positions (eg. those at the end of Chromosome 10 in D). “unmapped” indicates scaffolds of the *H. melpomene* reference genome that were not assigned to chromosomes in v1.1 of the genome assembly.

**Table 2.**
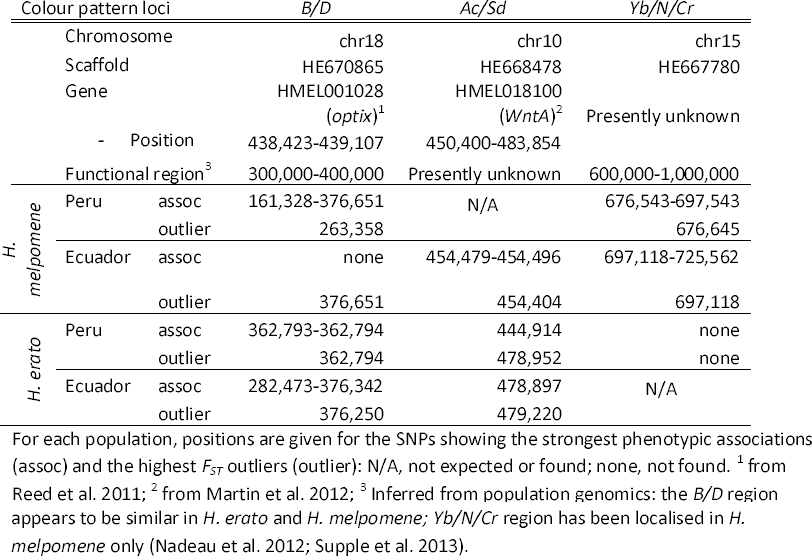
Positions of the strongest associations and outliers identified in genomic regions known to control colour pattern variation, in relation to known genes and functional regions.

Independent analyses were performed on both the reference alignments and *de novo* assemblies of the data. In all populations, more associated SNPs were identified in the alignments than in the *de novo* assemblies (Table 1, figure 5). We used blastn (Altschul et al. 1990) to place *de novo* contigs containing associated SNPs onto the *H. melpomene* genome, and most could be confidently assigned to a unique locus. There was almost no overlap in the particular SNPs detected in the differently assembled and aligned data sets (Figure 5), although in many cases the SNPs detected were in similar regions (Figures 3 and 4). There was some evidence for a higher false positive rate in the *de novo* data, as a higher proportion of associations were found scattered across the genome, away from the known colour pattern loci.

**Figure 4.**
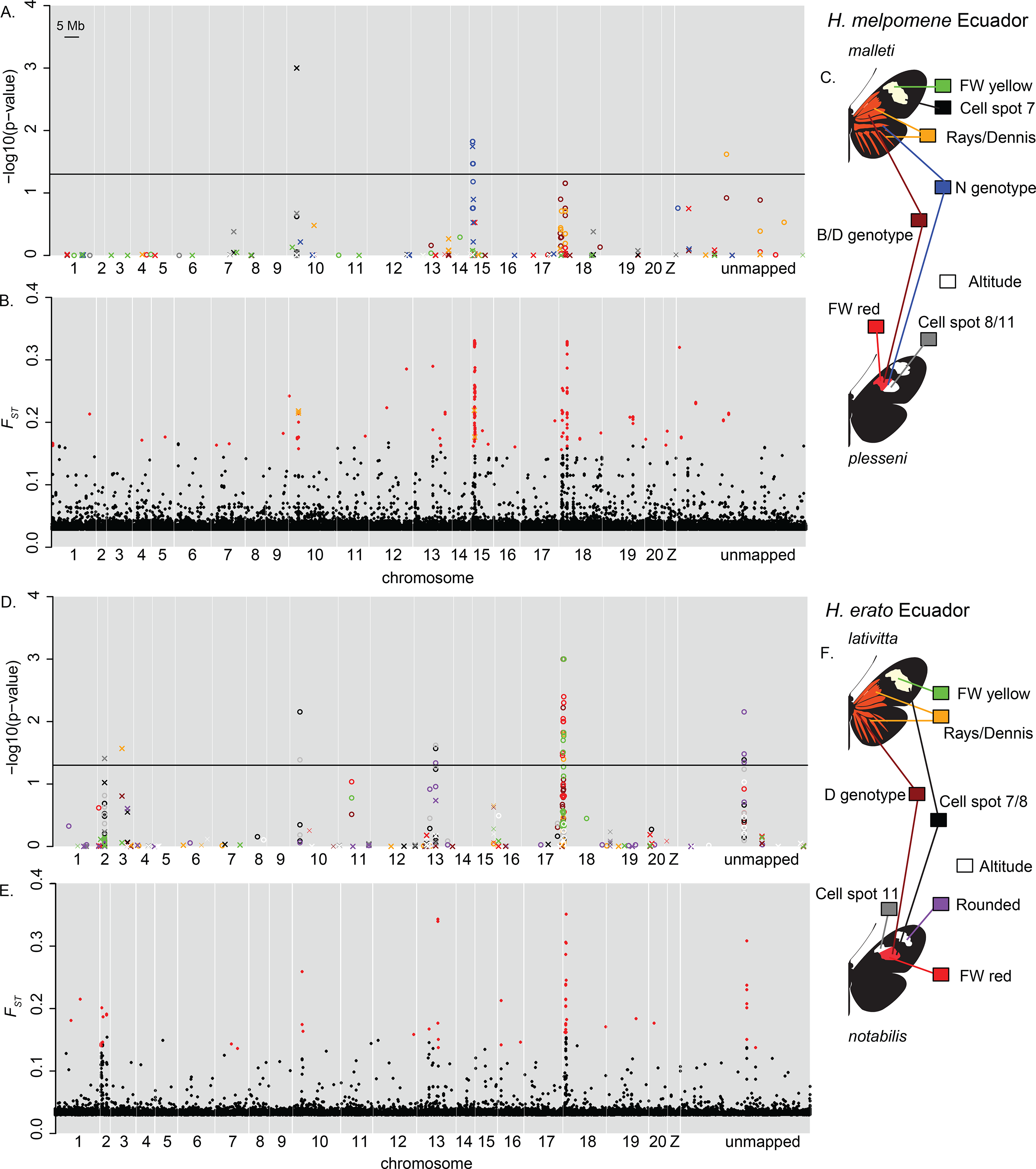
Association mapping (A and D) and outlier analysis (B and E) for *H. melpomene* (A, B, C) and *H. erato* (D, E, F) in Ecuador. See Figure 3 legend for further information.

**Figure 5.**
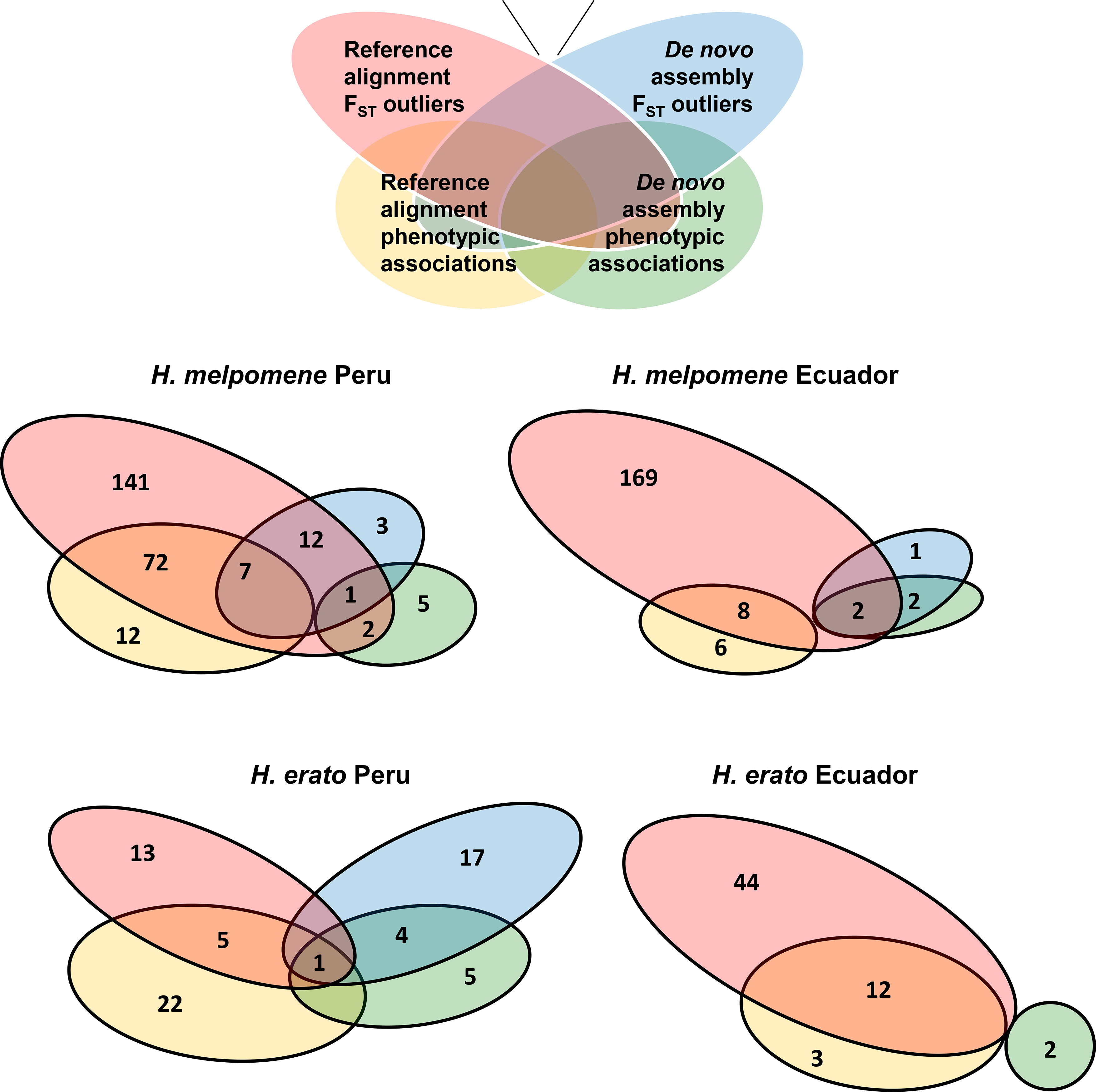
Venn diagrams of SNPs detected in the *de novo* assembled (orange and red) and reference aligned (blue and green) data by BayeScan outlier detection (red and blue) and association mapping (orange and green), for each of the four populations.

#### Red colour pattern elements and the B and D loci

Consistent with previous mapping studies based on crosses, red colour pattern variation in almost all populations was mapped to scaffold HE670865 on chromosome 18 (Baxter et al. 2008; Counterman et al. 2010; The *Heliconius* Genome Consortium 2012). The only exception was *H. melpomene* in Ecuador, where SNPs in this region did not reach significance (Figure 4A). This is likely to be due to the reduced sample size of this population (22) after removing the *H. timareta* individuals. Colour patterns were scored both as independent elements and also using known patterns of segregation to score the predicted genotype at the *B*/*D* locus (individuals with both red forewing and rays are predicted to be heterozygotes, as both traits are dominant). The latter scoring generally gave stronger associations (Figures 3 and 4), although both methods gave some significant associations for at least one of the traits. In all populations the strongest associations in this region were over 60 kb downstream of the *optix* gene that controls red colour pattern (Reed et al. 2011), but overlapped with the region identified in previous analyses as likely containing the functional regulatory variation (Nadeau et al. 2012; Supple et al. 2013)(Table 2, SI figure 4).

In the Peruvian *H. melpomene* population we also found weaker but significant associations with the *B*/*D* genotype on other linked chromosome 18 scaffolds, particularly HE671488. However, this scaffold was also associated with differences in altitude, which were stronger than the associations with colour (Figure 3A, Table 3, SI figure 4). This could suggest that this *B*/*D* linked region is responsible for ecological adaptation, although colour and altitude are strongly correlated so we do not have much power in the data set to separate the two. Both the Peruvian and Ecuadorian *H. melpomene* populations had a SNP at position 97 on an unmapped scaffold, HE670458, that was highly associated with rays (Table 3). This scaffold appears to consist largely of repetitive elements (BLAST hits match many other regions of the *H. melpomene* genome), suggesting that there may be a copy of a repetitive element that is associated with the presence of rays in both populations. All rayed individuals were heterozygous and all non-rayed individuals were homozygous at this SNP in both *H. melpomene* populations, which would be consistent with the presence of a unique copy of a repetitive element linked to the rayed allele. The existence of such a repetitive element is consistent with previous findings that repetitive elements are present in the region of highest divergence at the *B*/*D* locus (Nadeau et al. 2012; Papa et al. 2008).

**Table 3.**
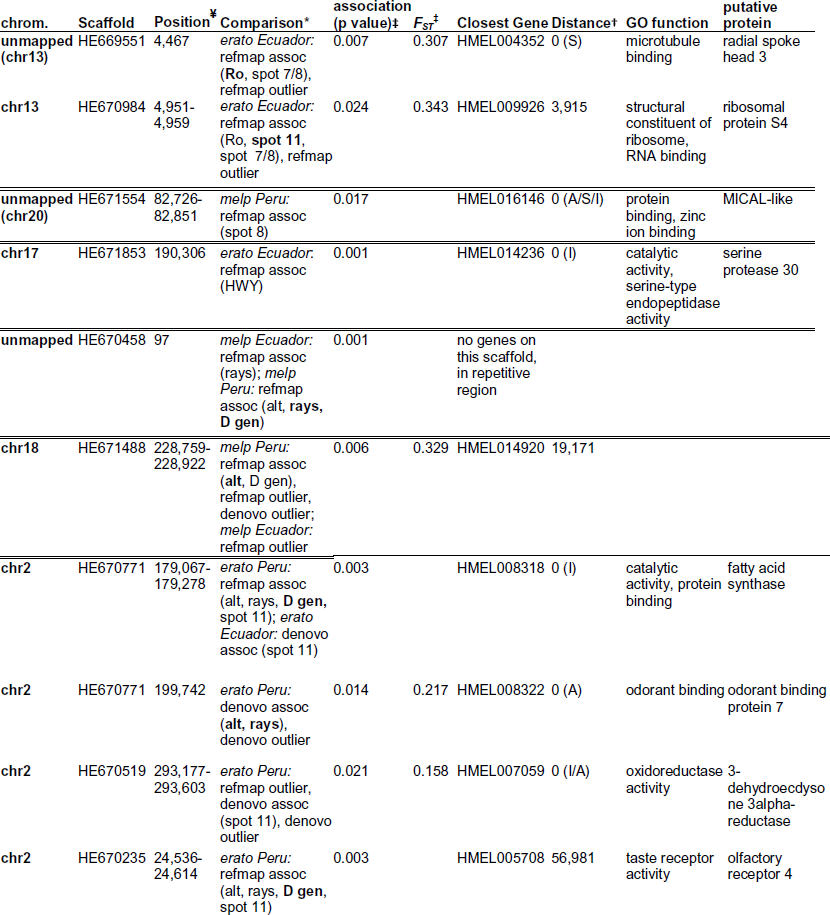
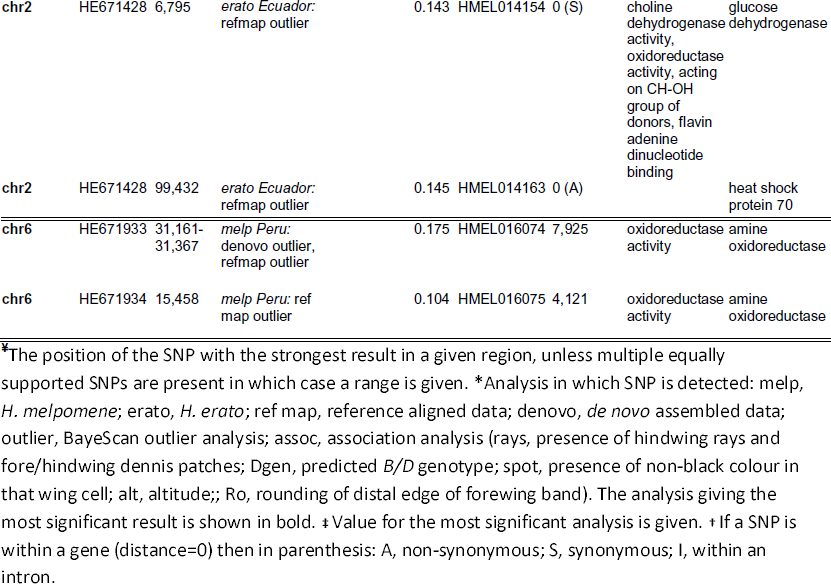
Novel genomic regions showing phenotypic associations or divergence outliers.

Surprisingly, in the Peruvian *H. erato* population the strongest associations with red colour pattern elements were not on chromosome 18, but at two scaffolds (HE670771 and HE670235) on chromosome 2 (Figure 3D, Table 3, SI figure 4). In addition, two SNPs significantly associated with rays and *D* genotype in the *de novo* assembled data of this population could not be confidently assigned to a position in the genome: One with a fairly weak top BLAST hit to the end of chromosome 10 and multiple equally strong hits elsewhere in the genome; the other had a top BLAST hit to an unmapped scaffold but additional good hits to two other unmapped scaffolds and chromosomes 2 and 19.

#### Yellow colour pattern elements and the Yb, N and Cr loci

In the Peruvian *H. melpomene* population the presence of the yellow hindwing bar and yellow in the forewing band both mapped to chromosome 15, with positions that were consistent with previous work on the *Yb* locus (Ferguson et al. 2010; Nadeau et al. 2012; The *Heliconius* Genome Consortium 2012)(Table 2, SI figure 4). Associations with altitude were also found at these associated SNPs but were weaker than the association with colour and so may simply be due to correlations between the altitude of the sampling site and colour pattern.

In the Peruvian *H. erato* population we did not recover the expected associations with the yellow hindwing bar, which is known to be controlled by the *Cr* locus on chromosome 15 (Joron et al. 2006; Counterman et al. 2010). Instead, the strongest association with this phenotype in the reference aligned data was found on chromosome 17 (Table 3). Moreover, we also identified significant associations with the yellow hindwing bar on chromosome 10 in both the reference aligned and *de novo* assembled data. In particular, one SNP on chromosome 10 was detected in both data sets on scaffold HE668478. These associations can be explained by the presence of *Sd* on this scaffold (Martin et al. 2012) (Table 2), which is known to influence the expression of the yellow hindwing bar, particularly in individuals that are heterozygous at the *Cr* locus (Mallet 1989). The *Cr* locus is not thought to control any aspects of phenotypic variation in Ecuadorian *H. erato* (Salazar 2012) and consistent with this expectation we did not detect any phenotypic associations in this region.

Significant associations with yellow in the forewing band were present in the *D* region in Ecuadorian *H. erato* (Figure 4D), consistent with the fact that the *D* locus controls both yellow and red colouration in the forewing band in *H. erato* (Sheppard et al. 1985, Salazar 2012, Papa et al. 2013). No significant associations were found with the presence of yellow colour in the forewing band in either Peruvian *H. erato* or Ecuadorian *H. melpomene*.

We scored the Ecuadorian *H. melpomene* individuals for their predicted genotype at the *N* locus (Figure 4C), which is known to control the amount and location of red in the forewing band and the length of the orange hindwing “dennis” bar (Salazar 2012). Despite the epistatic interaction between the *B*/*D* and the *N* loci, we could still score them independently. Associations with *N* were found to overlap the *Yb* region (Ferguson et al. 2010; Nadeau et al. 2012) on chromosome 15, scaffold HE667780 (Table 2, SI figure 4), in both the reference aligned and *de novo* assembled data (Figure 4A). These are the first genetic mapping results for the *N* locus and confirm previous studies’ findings based on segregation patterns that it is tightly linked to *Yb* (Sheppard et al. 1985; Mallet 1989).

#### Forewing band shape and the Ac, Sd and Ro loci

In Peruvian *H. melpomene*, the strongest associations with forewing band shape (cell spot 8 and cell spot 11) were on chromosome 18, within the *B*/*D* region (Figure 3A). This suggests that the *B*/*D* locus controls the shape as well as the colour of the forewing band in Peruvian *H. melpomene*. However, we did also find a cluster of 8 SNPs associated with band shape on an unmapped scaffold, HE671554. New mapping analyses suggest that this scaffold is on chromosome 20 (J Davey, Pers. Comm.) and therefore not linked to any previously described colour pattern controlling loci (Table 3).

In Peruvian *H. erato*, the SNP in the *Sd* region that was associated with the yellow hindwing bar also showed the expected association with forewing band shape in the *de novo* assembly but not the reference alignment. This SNP was just 5 kb upstream of the *WntA* gene (Table 2, Figure 3D, SI figure 4). Associations with forewing band shape were also found on chromosome 2 in this and the Ecuadorian *H. erato* populations (Figure 3D, Figure 4D, SI figure 4), in similar regions to those associated with red colour in Peruvian *H. erato* (Table 3).

In both species from the Ecuadorian hybrid zone, we found SNPs associated with forewing band shape (cell spot 7/8/11) within introns of the *WntA* gene (Figure 4, Table 2). In Ecuadorian *H. erato*, we also found two tightly linked SNPs on chromosome 13, scaffold HE670984, and three tightly linked SNPs on an unmapped scaffold, HE669551, which were associated with forewing band shape and also rounding of the band (Figure 4D). Rounding of the distal edge of the band in this population has previously been described as being under the control of the unmapped *Ro* locus (Sheppard et al. 1985; Salazar 2012). More recent mapping analysis suggests that HE669551 is within 1cM of HE670984 on chromosome 13 (J Davey, Pers. Comm.) so these associations are most likely due to a single locus on this chromosome. This is therefore the most likely genomic location of the *Ro* locus, representing the first time this locus has been mapped.

### *F_ST_* outlier detection

Outlier detection provides an alternative method for identification of loci under selection that does not depend on phenotypic association. BayeScan detected less than 0.06% of SNPs as outliers in each of the analyses (Table 1). In the *de novo* assembly, Peruvian hybrid zones showed a greater percentage of SNPs as outliers in both *H. erato* (0.059%) and *H. melpomene* (0.040%), with no outliers detected in Ecuadorian *H. erato* and only five in Ecuadorian *H. melpomene* (0.012%). The overall proportion of SNPs detected in the reference aligned data was similar. However, unlike the *de novo* assemblies, the proportions of outliers found in the reference alignments were more similar within each species than within each locality. Reference aligned data from *H. melpomene* contained approximately 0.025% outlier SNPs in both Peru and Ecuador, while reference aligned data from *H. erato* had 0.005% outliers in Peru and 0.017% outliers in Ecuador (Table 1). This would be consistent with some of the most rapidly diverging regions being lost in *H. erato* when aligned against the reference *H. melpomene* genome.

As suggested by results from the *de novo* assemblies, there do appear to be differences in population structure between the geographic regions that are consistent across both species. This is also reflected in the *F*_*st*_ distributions (from both alignment and assembly approaches), with both *H. erato* and *H. melpomene* having higher mean and background levels of *F_ST_* in Ecuador as compared to Peru (Table 1, Figure 2, SI figure 3). However, within both regions *H. melpomene* has a lower mean *F_ST_* than *H. erato*, which would be consistent with higher dispersal distances in *H. melpomene*, as previously suggested (Mallet et al. 1990). Similar outlier regions were detected by both the alignment and assembly approaches (Figures 3 and 4, B and E), although only Peruvian *H. melpomene* gave a good overlap in the specific SNPs detected (Figure 5). Some of the outlier contigs detected in Peruvian *H. erato* could not be positioned on the *H. melpomene* genome with confidence. In particular, one contig containing 4 outlier SNPs, had the highest BLAST homology to an unmapped scaffold (score = 87.7 bits) but similar hits (^∼^86 bits) to chromosomes 1 and 4. In addition, two of the contigs containing 3 outlier SNPs appeared to be linked to the *Sd* and *D* loci but were assigned with very low confidence (BLAST scores < 40 bits).

Overall, there was considerable overlap between the genomic regions containing outlier SNPs and those showing phenotypic associations (Figures 3 and 4), and to some extent in the specific SNPs, with the majority of phenotypically associated SNPs also being outliers (Figure 5). The exception to this general trend was the Peruvian *H. erato* population where a large proportion of the phenotypic associations were not strongly divergent between subspecies. In all populations over 50% of outlier SNPs were within 1Mb of a known colour pattern locus (including the newly identified *Ro* region; excluding these, 37.5% of outliers in Ecuadorian *H. erato* were within 1Mb of the *D* and *Sd* loci). The strongest outliers on chromosome 10 in the Ecuadorian populations and Peruvian *H. erato* were within intons of the *WntA* gene and the strongest outliers on the *B*/*D* scaffold were all 3’ of the *optix* gene (Table 2, SI figure 4).

In both *H. melpomene* populations there was a second strongly divergent region on chromosome 18 about 2Mb from the *B*/*D* region, which was not divergent in either of the *H. erato* populations (Figure 3B, SI figure 4). This is the same region on scaffold HE671488 that showed associations with colour pattern and altitude in the Peruvian *H. melpomene* population (Table 3). In the Peruvian *H. melpomene* population, we detected two clusters of outlier divergent SNPs on chromosome 6, which do not appear to be associated with colour pattern (Figure 3B, Table 3, SI figure 4). Outliers were also detected on chromosome 2 in both *H. erato* populations, some of which were in similar regions to those detected in the association mapping (Table 3, SI figure 4).

## DISCUSSION

It has long been recognised that convergent and parallel evolution provides a natural experimental system in which to study the predictability of adaptation (Stewart et al. 1987; Wood et al. 2005). This approach has come to the fore with the recent integration of molecular and phenotypic studies of adaptive traits (Stinchcombe and Hoekstra 2007; Nadeau and Jiggins 2010). Here, we have studied parallel divergent clines in two co-mimic species of butterflies, using RAD sequencing to generate an extensive data set covering 1-5% of the entire genome. Previous genomic studies of these species have sampled only a few individuals of divergent wing pattern races (Nadeau et al. 2012; The *Heliconius* Genome Consortium. 2012; Nadeau et al. 2013; Martin et al. 2013; Supple et al. 2013; Kronforst et al. 2013), while previous hybrid zone studies have yet to integrate next-generation sequencing approaches (Mallet and Barton 1989a; Salazar 2012; Baxter et al. 2010; Counterman et al. 2010). Here we have shown that association mapping in the hybrid zones can be used to find known loci, but also identify previously unmapped loci, such as *Ro* in *H. erato* and *N* in *H. melpomene*. Moreover, we conducted the first genome-wide scan for divergent loci and identify some that are not wing colour pattern related and so may have a role in other aspects of ecological divergence. With these data, we also identify a cryptic population of *H. timareta* in Ecuador and reveal parallel patterns of divergence between co-mimetic species.

### Comparison of de novo assembly and reference alignment of RAD data

Genome-wide association studies (GWAS) are now common in studies of admixed human populations (Visscher et al. 2012). Their use outside of model organisms has mostly been hampered by lack of reference genomes or methods for typing sufficient numbers of markers. However, these limitations are rapidly being eroded as the cost of sequencing decreases and more reference genomes become available. Furthermore, we have shown that alignment of reads to a fairly distantly related reference genome (^∼^15% divergent) can generate meaningful results. In the absence of a reference genome, *de novo* assembly also detects the same loci, but with somewhat reduced efficacy.

Alignment of sequence reads to the reference genome produced data for more sites, even in the more distantly related species, *H. erato*. One drawback of the STACKS pipeline that we used for *de novo* assembly of the reads is that it does not assemble and call sequence variants in the paired end reads. Hence the available sequence for analysis is almost double in the reference alignments as compared to the de novo assembly. However, it also seems that data was lost in the to *de novo* assembly due to divergent alleles not being assembled together. This may have had a larger influence on the *H. erato* assemblies as this species harbours greater genetic diversity than *H. melpomene* (Hines et al. 2011) and so explain why a much lower proportion of the *de novo* assembled contigs were present across multiple individuals in *H. erato* (Table 1). We also found a higher proportion of variable sites in the reference alignments as compared to the *de novo* assemblies. This may again be due to poor assembly of the *de novo* contigs, but it could also represent genetic variability contained in the paired-end reads. It is possible that paired end reads might be located in more variable regions, particularly if restriction-site associated reads were biased towards more conserved regions (The *Heliconius* Genome Consortium 2012).

The larger number of SNPs in the reference alignments resulted in larger numbers of outlier and associated SNPs being detected, most of which cluster in the expected genomic regions. Moreover, there appears to be a higher false positive rate in the association mapping using the *de novo* assembled data. The most likely explanation for this result is that the smaller number of SNPs generated from the *de novo* assembly gave less power to correct for underlying population structure. Nevertheless, many of the expected associations and outlier regions were detected in the *de novo* assembled data. The results from assembly and alignment approaches are more concordant in *H. melpomene* than *H. erato* particularly at the level of individual SNPs (Figure 5). This is very likely due to the fact that the *H. melpomene* reference genome was used to generate the sequence alignments in both species. In addition, the lower within-population diversity in *H. melpomene*, may also have led to improved *de novo* assemblies in this species.

Overall, our results suggest that detection of loci underlying adaptive change is likely to be more effective where reads can be mapped to a reference genome. The *de novo* approach could, and no doubt will, be improved by developing methods that allow paired end reads to be incorporated into the SNP typing pipeline (Baxter et al. 2011). This would not only allow a higher density of SNPs to be detected but could also improve alignment of divergent alleles. In the mean time, one approach that has been used in other studies is to *de novo* assemble RAD-seq reads to generate a consensus reference that the reads are then mapped back to for SNP calling (Keller et al. 2013).

### Association mapping across hybrid zones is a rapid way of detecting loci underlying phenotypic differences

We have successfully used association mapping in hybrid zone individuals to identify virtually all of the genomic regions known to control colour pattern in these populations (Reed et al. 2011, Martin et al. 2012; Nadeau et al. 2012; Supple et al. 2013). It has commonly been supposed that large sample sizes will be necessary in order identify genes in wild populations. Here we have confirmed recent theoretical predictions from simulated data (Crawford and Nielsen 2013), that for large effect adaptive loci, even small sample sizes can be highly effective in identification of narrow genomic regions underlying adaptive traits. We also confirm the prediction that in populations with low background levels of divergence both divergence outlier and association mapping approaches are effective in detecting regions under divergent selection. In our study, association mapping has the added benefit of identifying the phenotypic effects of the selected loci. One anticipated pitfall of this method was that many of the phenotypes covary across the hybrid zone. However, it appears that with just 10 individuals with admixed phenotypes we can disassociate most of the variation and thus find distinct genetic associations for known loci. This therefore gives us some confidence that the novel associations that we have detected are real and not due to covariation with other phenotypes.

In Ecuador we intentionally sampled from sites at the edges of the hybrid zone where both pure and hybrid individuals were present, as we anticipated that individuals from these sites would have the highest levels of admixture between selected alleles. This may explain the clearer patterns observed in *H. erato* in Ecuador as compared to Peru (Figures 3D and 4D). In Peruvian *H. erato* we also find several genomic regions showing phenotypic associations that are not divergence outliers, which may suggest that these are likely to be false positives. However, the less clear signal in Peruvian *H. erato* could also be due to the reduced sample size in this population (27 individuals). Certainly, the reduced number of Ecuadorian *H. melpomene* individuals (22) seems to have greatly reduced the power of the association mapping to detect significant associations (Figure 4B). The lack of any signal at the *Cr* locus in Peruvian *H. erato* is still surprising, and may be because the *Cr* associated region is very narrow. Linkage disequilibrium breaks down rapidly in *H. erato* (Counterman et al. 2010) and even though there are >700 SNPs in *H. erato* within the region that contains hindwing bar associated SNPs in *H. melpomene*, this may not be enough to identify the loci responsible for this phenotypic variation. Nevertheless, contrary to previous suggestions (Kronforst et al. 2013), the density of RAD markers we have obtained was sufficient to identify many other narrow divergent genomic regions.

Although we have clearly demonstrated the utility of this approach for association mapping, it should be noted that scoring of some phenotypes was informed by previous crossing experiments. For example, the *N* locus in Ecuadorian *H. melpomene* was scored taking into account the genetic background at *B*/*D* (Salazar 2012), and the scoring of the predicted genotype at the *B*/*D* locus yielded stronger associations than scoring of individual colour pattern elements. Nonetheless, scoring based purely on phenotypic variation did successfully identify colour pattern loci in several cases (eg. *Ro, Ac/Sd* and *Yb)*. Overall, the prospects for mapping individual phenotypic components and identifying epistatic relationships without prior knowledge are considerable, especially with larger sample sizes.

The possibility of using hybrid zones for association mapping has long been recognised (Kocher and Sage 1986) but few studies have successfully applied this technique. Studies in younger hybrid zones, for example *Helianthus* sunflowers, have found that linkage disequilibrium between unlinked genomic regions in early generation hybrids can produce spurious associations (Rieseberg and Buerkle 2002). *Heliconius* hybrid zones seem ideal in this regard because they appear to be fairly ancient and close to linkage equilibrium. However, it seems likely that many other suitable systems do exist for this type of approach (Lexer et al. 2006; Crawford and Nielsen 2013). An additional benefit of the *Heliconius* system is that much of the phenotypic variation is controlled by major effect loci, which can be detected with small sample sizes. Much larger sample sizes would be required in order to detect minor effect loci (Beavis 1997). Nevertheless, adaptive phenotypes appear to frequently arise from major effect loci (Orr 2005; Nadeau and Jiggins 2010) and so there may be many situations in which the use of very large sample sizes is unnecessary. In addition, by incorporating methods that use a probabilistic framework to infer allele frequencies in low coverage sequencing data (Gompert and Buerkle 2011) it should be feasible to sequence large enough samples for analysis of quantitative traits.

### Identification of a novel colour pattern locus

Our association mapping results have robustly identified the *H. erato Ro* locus, that controls the shape of the distal edge of the forewing band, as being on chromosome 13 near gene HE669551. This gene has a predicted Gene Ontology (GO) molecular function of microtubule binding and is similar to other insect Radial Spoke Head 3 proteins, which are components of the cilia (Avidor-Reiss et al. 2004). It is therefore not an obvious candidate for control of colour pattern, so may simply be linked to the causative site. Our results are contrary to the suggestion of a recently published QTL study that *Ro* may be linked to *Sd* (Papa et al. 2013). However, that study also identified a major unlinked QTL for forewing band shape, that could not be assigned to a *H. melpomene* chromosome and so may be homologous to the locus we detected here. Furthermore, a QTL for several aspects of forewing band shape and size, including the shape of the distal edge, has previously been identified in *H. melpomene* on chromosome 13 (Baxter et al. 2008). This was located to a fairly broad region but its positioning is consistent with our results for the *Ro* locus in *H. erato*. It therefore seems likely that we have identified a new wing patterning locus that is homologous in *H. melpomene* and *H. erato*.

### Ecological selection across the hybrid zones

Our results support previous assertions that selection acting on colour pattern is the most important factor in maintaining these hybrid zones (Mallet and Barton 1989a; Baxter et al. 2010; Counterman et al. 2010; Nadeau et al. 2012; Supple et al. 2013). The most divergent genomic regions correspond to colour pattern controlling loci and at least half of all divergence outliers are in these regions. Nevertheless, some divergent regions do not seem to correspond to colour pattern loci, and could be candidates for adaptation to other ecological factors. The best candidates appear to be the regions on chromosome 2 in *H. erato* and chromosome 6 in *H. melpomene*. The regions on chromosome 2 in the Peruvian *H. erato* population are also associated with colour pattern, but such association could be due to the high covariation of colour pattern and sampling location in this population. These regions overlap with predicted genes, including basic metabolic genes and a heat shock protein (Table 3), which could be candidates for adaptation to different temperature regimes. Chemosensory genes were also detected on chromosome 2, and could be candidates for divergent mate preference or host plant adaptation (Briscoe et al. 2013). However, no differences in host plant preference have been observed in Peru where these outliers were detected, and mating within the hybrid zone appears to the random (Mallet and Barton 1989a), although marginal differences in mate preference have been observed in *H. melpomene* (Merrill, Gompert, et al. 2011).

There appear to be multiple dispersed divergent regions on chromosome 2 in *H. erato* (SI figure 4). These could be evidence of divergence hitchhiking, whereby new mutations that cause differential fitness are more likely to be fixed by selection if they arise close to other loci already under divergent selection. This could lead to clustering of divergently selected loci in the genome (Via 2012; Feder et al. 2012). The same process could also have led to additional loci under divergent selection to arise in linkage with the colour pattern loci. This could explain the second divergent and altitude associated region on chromosome 18 (linked to the *B*/*D* locus) in *H. melpomene* (Table 3, SI figure 4). One possibility is that this could be the *B*/*D* linked mate preference locus that has previously been identified (Merrill, Van Schooten, et al. 2011), although it is not clear if the mate preference locus is an additional linked locus or a pleiotropic effect of the wing colour locus itself. It is also possible that these apparently distinct but linked divergent regions could simply reflect the heterogeneous nature of *F_ST_* resulting from strong divergent selection acting on a single locus combined with other background and neutral processes (Charlesworth et al. 1997). A broad region of divergence around the *B*/*D* locus in *H. melpomene* would fit with other suggestions that it has undergone stronger or more recent selection than other colour pattern loci (Nadeau et al. 2012). The *D* region in *H. erato* does not appear to be extended in the same way as in *H. melpomene* (SI figure 4), suggesting that either the architecture or the selective history of this region is different between these species.

### Comparison of the genomic architecture of divergence between convergent species

One interesting question that can be addressed with our results is the extent to which species undergoing parallel divergence will show parallel patterns at the genomic level. In order to address this we first need to know whether the species really have undergone parallel divergence, *i.e*. that both the phenotypic start and end points have been similar. Several previous studies have suggested that this is not the case and that *H. erato* diverged earlier and followed a different trajectory compared to *H. melpomene* (Quek et al. 2010; Brower 1996; Flanagan et al. 2004). However, our phylogenetic results are more consistent with a recent analysis suggesting that the two species do appear to have undergone co-divergence in multiple populations across their range (Cuthill and Charleston 2012). Our results are based on significantly more data than any of the previous analyses (>5Mb in *H. melpomene* and >1Mb in *H. erato*), and should produce a better signal for phylogenetic analysis as compared to AFLPs used previously (Quek et al. 2010). Although the striking similarities in tree topology do seem to support the co-divergence hypothesis, alignment to a reference genome means that the evolutionary rates in our data for *H. erato* and *H. melpomene* are not directly comparable. In addition to the phylogenetic signal, our data also suggested similar patterns of population structure between species in each of the regions, with higher background divergence levels in Ecuador as compared to Peru (Figure 2, Table 1, SI figure 3). This is despite the fact that average distance between sampling locations of “pure” subspecies individuals was similar for both hybrid zones (^∼^56 km in Ecuador and 58-60 km in Peru).

Although some loci show parallel divergence in both species (*B*/*D* in Peru; *B*/*D* and *Ac/Sd* in Ecuador), there is surprisingly little similarity in which other loci are divergent between subspecies within each species. This is contrary to the general perception that there are strong genetic parallels in this system (Joron et al. 2006; Baxter et al. 2008; Supple et al. 2013; Papa et al. 2008). Some of these differences were known previously, for example, that in Peru the *Sd/Ac* locus controls band shape variation in *H. erato* but not in *H. melpomene* (Mallet 1989). Our results extend this further through the identification of the *Ro* locus on chromosome 13 in Ecuadorian *H. erato*, which is not divergent in its co-mimic *H. melpomene*, and the identification of divergent regions of chromosome 2 in *H. erato* and chromosome 6 in Peruvian *H. melpomene*.

In general, it seems that although the same colour pattern loci are present in both species (Joron et al. 2006; Baxter et al. 2008; Martin et al. 2012) they are being used in different ways and combinations in order to produce convergent phenotypes. This is particularly surprising given the pattern of co-divergence observed in the phylogeny, which would appear to suggest that similar colour patterns have arisen at a similar time and from similar ancestral forms in both species. Nonetheless, the apparent pattern of co-divergence could simply reflect more recent patterns of gene flow between geographically proximate populations in both species. This has recently been highlighted by studies showing that patterns of divergence at colour pattern controlling loci can be very different to those found at the rest of the genome (Hines et al. 2011; Pardo-Diaz et al. 2012; The *Heliconius* Genome Consortium 2012; Supple et al. 2013). Therefore, the differences that we observe in the use of particular loci in the two species could reflect different mimetic histories that will only be resolved by studies of the evolutionary history of particular loci.

### Discovery of a new cryptic *H. timareta* population

An unexpected finding of our study was the discovery of a previously undescribed population of H. *timareta*, which appears phenotypically virtually indistinguishable from *H. melpomene malleti* in Ecuador but is clearly genetically distinct (Figure 1, SI figures 1 and 2). *H. timareta florericia* is a *malleti* like population that has previously been described in Colombia and also co-occurs with *H. melpomene malleti*. In that population the length of the red line on the anterior edge of the ventral forewing was diagnostic (Giraldo et al. 2008). This character was not diagnostic in our genotyped individuals, with overlapping length distributions between the species (data not shown). We noted a tendency towards *H. timareta* having a shorter line on average, but given the small sample sizes in the current study, this remains to be confirmed.

A polymorphic high altitude population of *H. timareta (H. timareta timareta)* also occurs in this area of Ecuador, overlapping in distribution with *H. melpomene plesseni*. The polymorphism in this population has been somewhat of a puzzle as none of the forms mimic other co-occurring butterflies (Mallet 1999). Our finding of a new *H. timareta* population may help to explain the polymorphism in *H. timareta timareta*, if it is being generated in part by gene flow from this newly identified population.

The *H. timareta* radiation has only been recognised in the last 10 years (Giraldo et al. 2008; Jiggins 2008, Merot et al., 2013). The *H. timareta* individuals in our study were collected from sites at 824 m and 376 m. They appear to be fairly common at low altitude as four out of the five individuals sequenced from the site at 376 m were *H. timareta*. In a large dataset compiled by Rosser et al. (2012) containing 232 *H. timareta* individuals from all known populations (including *H. tristero*, now thought to be a subspecies of *H. timareta*), the lowest sampling location is around 600 m, with 95% of individuals occurring over 800 m. Therefore, the population of *H. timareta* that we have discovered occurs below the usual altitudinal range of *H. timareta*. This extends the possible range of this species and suggests that the overlap in distribution of *H. timareta* and *H. melpomene* is greater than previously considered.

### Conclusions

We have demonstrated that high resolution genome scans using admixed individuals from hybrid zones can be used to identify loci underlying phenotypic variation. Only a small proportion of the genome (about 0.025%) is strongly differentiated between subspecies and most of this can be explained by divergence at loci controlling colour pattern. This is consistent with previous studies based on smaller numbers of markers (Turner 1979; Baxter et al. 2010, Counterman et al. 2010, Nadeau et al. 2012) and suggests that the hybrid zones are ancient or have formed in primary contact, and are maintained by strong selection on colour pattern (Mallet and Barton 1989a, Mallet 2010). However, we also find, for the first time, some divergent loci that do not appear to be associated with colour pattern, suggesting that there may be other differences between subspecies. This could explain why several *Heliconius* hybrid zones occur across ecological gradients (Benson 1982), if they are coupled with extrinsic selection acting on other loci in the genome (Bierne et al. 2010). However, this needs to be confirmed with detailed phenotypic analyses of the subspecies to identify whether differences are present that could be explained by ecological adaptation. In general we find that, although some loci are divergent in all populations, the genomic pattern of divergence between co-mimetic species is not particularly similar, suggesting that the level of parallel genetic evolution between *H. erato* and *H. melpomene* is in fact quite low, despite parallel phylogenetic patterns of divergence. Finally, our analysis shows that alignment to a distantly related reference genome can improve analyses over a *de novo* assembly of the data.

## METHODS

### Samples and sequencing

30 *H. erato* and 30 *H. melpomene* individuals were selected from a larger sample taken from the hybrid zone region in Peru. Similarly, 30 *H. erato* and 30 *H. melpomene* were also selected from a larger study of a subspecies hybrid zone in Ecuador (Salazar 2012). Each set of 30 samples comprised 10 pure forms of each subspecies and 10 hybrids (based on colour pattern). See Figure 2 and SI table 2 for further details of the samples and locations.

RAD sequencing libraries were prepared using previously described methodologies (The *Heliconius* Genome Consortium 2012; Baird et al. 2008; Baxter et al. 2011). Briefly, DNA was digested with the restriction enzyme Pstl prior to ligation of P1 sequencing adaptors with five-base molecular identifiers (see SI table 2 for MIDs used). We then pooled samples into groups of 6 before shearing, ligation of P2 adaptors, amplification and fragment size selection (300–600 bp). Libraries were then further pooled such that 30 individuals were sequenced on each lane of an Illumina HiSeq2000 sequencer to obtain 150 base paired-end sequences. We obtained an average of 374M sequence pairs from each lane. Following sequencing, three of the *H. erato* individuals from Peru were found to have been incorrectly assigned to this species and were excluded from all further analyses.

In order to compare patterns of phylogenetic divergence of the focal subspecies, we also used sequence data from additional subspecies and closely related species in each group. Two individuals each from 6 additional *H. erato* populations and the closely related *H. himera* were also Pstl RAD sequenced with 5 individuals pooled per lane of Illumina GAIIx (100 base paired-end sequencing). These sequences were obtained in the same run as a comparable set of individuals from the *H. melpomene* clade, which have been used in previous analyses (The *Heliconius* Genome Consortium 2012; Nadeau et al. 2013, European Nucleotide Archive, Accession ERP000991). We also obtained whole-genome shotgun sequence data from an outgroup species, *H. clysonimus*, which was sequenced on a fifth of a HiSeq2000 lane, giving 53.5M 100 base read pairs for this individual.

### Alignment to reference genome

We separated paired-end reads by MID using the RADpools script in the RADtools (vl.2.4) package (Baxter et al. 2011), which also filters based on the presence of the restriction enzyme cut site, using the option to allow one mismatch within the MID. Reads from each individual were then aligned to the *H. melpomene* reference genome (The *Heliconius* Genome Consortium 2012) using Stampy vl.0.17 (Lunter and Goodson 2011), with default parameters except substitution rate, which was set to 0.03 for alignments of *H. melpomene* and 0.10 for alignments of *H. erato*.

We then realigned indels and called genotypes using the Genome Analysis Tool Kit (GATK) vl.6.7 (DePristo et al. 2011), emitting all confident sites (those with quality ≥30). This was first run on each set of 30 (or 27) individuals from each population group. These genotype calls were used for analyses of genetic variation within each of the groups, including outlier detection, association mapping and analyses of subpopulation structure. In addition, genotype calling was also performed on a combined dataset of all *H. melpomene* and outgroup taxa *(H. timareta, H. cydno* and *H. hecale)* as well as a combined set of all *H. erato* and its outgroups *(H. himera* and *H. cylsonimus)*. These genotype calls were used for the phylogenetic analyses and broader analyses of genetic structure. For all downstream analyses, calls were further filtered to only accept those based on a minimum depth of five reads and minimum genotype and mapping qualities (GQ and MQ) of 30 for *H. melpomene* and 20 for *H. erato*.

### *De novo* assembly

We quality-filtered the single-end raw sequence data and separated sequences by MID with the process_radtags program within Stacks (Catchen et al. 2011). This program corrects single errors in the MID or restriction site and then checks quality score using a sliding window across 15% the length of the read. We discarded sequences with a raw phred score below 10, removed reads with uncalled bases or low quality scores, and trimmed reads to 100 bases to eliminate potential sequencing error occurring at ends of reads. Table 1 shows the mean read numbers per individual obtained after filtering. For each population group, we assembled loci *de novo* using the denovo_map.sh pipeline in Stacks (Catchen et al. 2011). We set the minimum depth of coverage (m) to 6, allowed 4 mismatches both in creating individual stacks (M) and in secondary reads (N), and removed or separated highly repetitive RadTags. Due to the high level of polymorphism in our dataset, we used these parameters to minimize the exclusion of interesting loci with high variability between populations. *De novo* assembly was conducted both including (for association mapping) and excluding (for bayescan outlier detection) hybrid individuals in the analysis. Individuals from Ecuador that were identified as being *H. timareta* were excluded.

### Phylogenetics and analysis of population structure

Only the reference aligned data were used for phylogenetics and Structure analyses. We used custom scripts to convert from vcf to Phylip format and to filter sites with a minimum of 95% of individuals with confident calls. Maximum likelihood phylogenies were constructed in PhyML (Guindon and Gascuel 2003) with a GTR model using the resulting 5,737,351 sites (including invariant sites) for the *H. melpomene* group and 1,693,024 sites for the *H. erato* group. Approximate likelihood branch supports were calculated within the program.

Population structure within and across each of the hybrid zones was analysed using the program Structure v2.3 (Pritchard et al. 2000). We prepared input files using custom scripts, and only sites with 100% of individuals present for *H. melpomene* populations or at least 75% of individuals present for *H. erato* populations and with a minor allele frequency of at least 20% were retained. This reduced the number of sampled sites, keeping just the most informative ones, for easier handling by the program. Initial short runs (10^3^ burn-in, 10^3^ data collection, K = 1) were used to estimate the allele frequency distribution parameter λ. We then ran longer clustering runs (10^4^ burn-in, 10^4^ data collection) with the obtained values of λ for each of the four population groups for K = 1-3. For *H. melpomene* in Ecuador the analysis was first run with all individuals included and then excluding the individuals identified as being *H. timareta*.

We also performed principle components analysis of the genetic variation in each population group. This was done with the “cmdscale” command in R (R Development Core Team 2011), using genetic distance matrices calculated as 0.5-ibs, where ibs was the identity by sequence matrix calculated in GenABEL (see below). As further confirmation that some of the *H. melpomene* individuals sampled in Ecuador were in fact cryptic *H. timareta*, we also performed principle components analysis on the combined *H. melpomene* and outgroup data set. We also ran principle components analysis on the *de novo* assembled data for each population group, to test whether both methods were detecting similar underlying patterns of genetic variation.

In order to compare our newly identified *H. timareta* individuals to other populations, we Sanger sequenced a 745 bp region of mitochondrial COI that overlapped with the regions sequenced in previous studies (Giraldo et al. 2008; Mérot et al. 2013). This was PCR amplified as in Mérot et al. (2013) with primers “Jerry” and “Patlep” and directly sequenced with “Patlep”. These sequences were then aligned with those available on Genbank and a maximum likelihood phylogeny was constructed in PhyML (Guindon and Gascuel 2003) with a GTR model and 1000 bootstrap replicates.

### Association mapping of loci controlling colour pattern variation

We scored components of phenotypic variation that segregate across each of the hybrid zones. The scored phenotypes are shown in Figure 3 (for Peru) and Figure 4 (for Ecuador) and listed in full in SI Table 3. These were scored mostly as binomial (1,0) traits, but in some cases intermediates were also scored (as 0.5). The width and shape of the forewing band was scored based on whether it extended into each of the wing “cells”, demarcated by the major wing veins (as shown in SI figure 5). In Peruvian populations, the size and shape of the forewing band was measured as two components (Figure 3C/F) that extend the band distally (cell spot 8) and proximally (cell spot 11). In Ecuador, three aspects of band shape were scored: cells 8 and 11, which make up the proximal spot in *H. m. plesseni* and *H. e. notabilis*, and cell 7, which pushes the band towards the wing margin in *H. m. malleti* and *H. e. lativitta* (Figure 4C/F). In our sample of *H. melpomene* the presence of cell spots 8 and 11 were perfectly correlated, whereas in *H. erato* the presence of cell spot 7 was perfectly correlated with the absence of cell spot 8. In addition, individuals were also scored for their predicted genotypes at major loci described previously (with predicted heterozygotes scored as 0.5) (Sheppard et al. 1985; J. Mallet 1989) and the altitude at which they were collected was included as a continuous phenotypic trait.

We performed association mapping using the R Package GenABEL ν 1.7-4 (Aulchenko et al. 2007). This was performed on both the *de novo* assembled and the reference aligned data with a custom script used to convert both from vcf to Illumina SNP format. Individuals identified as being H. *timareta* were excluded. Filtering was performed within the program to remove sites with >30% missing data and with a minor allele frequency of <3%.

For each population, an analysis of the hind-wing ray phenotype using the reference mapped data was first performed using three methods: a straight score test (qtscore), a score test with the first three principle components of genetic variation (calculated as described above) as covariates, and an Eigenstrat analysis (egscore, Price et al. 2006). The presence of genetic stratification and the ability of these methods to correct for this was analysed by comparing the inflation factor, λ. In all cases the analyses incorporating population stratification did not give a reduced value of λ and so were not used for subsequent analyses. As our samples were from hybrid zones with >60% of the samples having extreme values of all scored phenotypes, we would expect similar levels of stratification for all phenotypes, so this test for stratification was not repeated for all phenotypes.

We therefore performed score tests for all scored phenotypes across all population groups. Genome-wide significance was determined empirically from 1000 resampling replicates and corrected for population structure using the test specific λ.

### BayeScan analysis to identify loci under selection

We used the program BayeScan v2.1 (Foll and Gaggiotti 2008) to look for loci with outlier *F_ST_* values between “pure” individuals of each subspecies type (based on wing colour pattern) in each population group. Exclusion of the *H. timareta* individuals meant that only three pure *H. melpomene malleti* individuals remained. Therefore, for the purpose of this analysis of *H. melpomene* in Ecuador, the two hybrid individuals closest to the *H. m. malleti* side of the hybrid zone (Figure 2), which also had the most *H. m. malleti* like phenotypes, were included as *H. m. malleti*.

The program was run with the prior odds for the neutral model (pr_odds) set to 10 and outlier loci were detected with a false discover rate (FDR) of 0.05. We ran this analysis using both the *de novo* assembled and the reference aligned data. Custom scripts were used to convert these to the correct input format. For both analyses, sites were only kept if at least 75% of individuals were sampled for both subspecies in a given comparison.

## DATA ACCESS

DNA sequence reads have been submitted to the European Nucleotide Archive, Accession PRJEB4669. COI sequences have been deposited in EMBL-Bank, accessions HG710096 - HG710125. Custom scripts and wing images are available on request from the authors.

## ACKNOWLEDGEMENTS

We thank Simon Baxter, Doug Turnbull and William Cresko for their help and advice with RAD library preparation and sequencing. Sequencing was performed at the University of Oregon, Genomics Core Facility and The Gene Pool genomics facility in the University of Edinburgh. We would also like to thank Julian Catchen for his help with Stacks. We thank the governments of Peru and Ecuador for their permission to collect and export specimens. Santiago Villamarín from the Museo Ecuatoriano de Ciencias Naturales provided institutional support in Ecuador. We also thank Ismael Aldás, Carlos Robalino and Patricia Salazar for their assistance with fieldwork. Joanna Riley assisted with DNA extractions. John Davey gave us access to his unpublished mapping results. This project was supported by research grants NSF-CREST #0206200, NSF-DEB-1257839 and NSF-IOS 1305686. NJN was funded by a Leverhulme Trust award to CDJ, MR was funded through the Ford Foundation Postdoctoral Fellowship Program administered by the National Academies, HOZ and JAM were partially supported by NIH-NIGMS INBRE award P20GM103475 and NSF-EPSCoR award 1002410.

